# Interpreting Molecular Dynamics Forces as Deep Learning Gradients Improves Quality Of Predicted Protein Structures

**DOI:** 10.1101/2023.10.03.560775

**Authors:** Jonathan Edward King, David Ryan Koes

**Affiliations:** Joint PhD Program in Computational Biology, Carnegie Mellon University-University of Pittsburgh, Pittsburgh, PA 15213; Computational & Systems Biology, University of Pittsburgh, Pittsburgh, PA 15213

## Abstract

Protein structure predictions from deep learning models like AlphaFold2, despite their remarkable accuracy, are likely insufficient for direct use in downstream tasks like molecular docking. The functionality of such models could be improved with a combination of increased accuracy and physical intuition. We propose a new method to train deep learning protein structure prediction models using molecular dynamics force fields to work toward these goals. Our custom PyTorch loss function, OpenMM-Loss, represents the potential energy of a predicted structure. OpenMM-Loss can be applied to any all-atom representation of protein structure capable of mapping into our software package, SidechainNet. We demonstrate our method’s efficacy by finetuning OpenFold. We show that subsequently predicted protein structures, both before and after a relaxation procedure, exhibit comparable accuracy while displaying lower potential energy and improved structural quality as assessed by MolProbity metrics.

**SIGNIFICANCE:** We propose a novel framework to directly incorporate forces from molecular dynamics as gradients for training deep learning models like AlphaFold2. We implement our method as a PyTorch loss function, OpenMM-Loss, which frames the potential energy of predicted protein structures as a minimization objective. When applied to OpenFold, our method demonstrates improved structural quality, lower potential energy, and comparable accuracy relative to the OpenFold baseline. Our framework may enhance the ability of deep learning models to recapitulate fundamental biophysical principles, reducing the number of structural irregularities in their predictions and paving the way for more effective downstream applications.

## INTRODUCTION

Deep learning (DL) methods for protein structure prediction, particularly AlphaFold2 (AF2) (1) and AF2-like systems (2–4), have proven remarkably accurate. However, barriers to the practical use of AF2 predictions remain. Some of AF2’s shortcomings are due to limitations of its design, while others are simply areas where accuracy could be improved. For instance, Jumper et al. (1) trained AF2 on protein structures from the Protein Data Bank (PDB) (5). By virtue of their presence in the PDB, many of these proteins formed stable crystal structures in the lab. Such structures may not adequately reflect their *in vivo* conformations, which may exhibit higher energy under crystallization conditions. Another fundamental limitation of AF2 is its inability to simultaneously model proteins and their ligands. Without knowledge of protein-ligand interactions, AF2 cannot reliably predict *apo* or *holo* protein states, making drug development with these structures more challenging.

AF2 also likely needs higher accuracy for downstream tasks like docking. Several studies (6–8) have demonstrated that AF2-predicted structures are surprisingly non-performant compared to crystal structures used for docking. Despite a relatively high level of accuracy of the protein backbone and binding sites, subtle differences between the prediction and crystal structure cause significantly different results when docking molecules. Sidechain atom accuracy is a likely culprit since differences in sidechain atom placement or orientation may dramatically alter the binding pocket shape or chemical environment.

In addition to working toward more accurate models, the research community has expressed interest in developing models capable of understanding and recapitulating fundamental physical principles. Models that leverage existing scientific knowledge could free up computational resources to focus on subproblems better suited for machine learning instead of relearning principles taught in high school chemistry and mathematics. Such a model could, in principle, better guide drug discovery campaigns since it may better understand the chemical interactions between a protein and a ligand.

Researchers in several fields have tried utilizing more complex and physically relevant input representations and model architectures or even empirical data to improve prediction quality. For example, data representation in cheminformatics has shifted from SMILES (9) (string-based) representations of chemical structures to other representations like graphs (10, 11), point clouds (12), or 3D voxels (13) that more closely resemble the physical reality of these systems. One DL method for computational drug discovery utilizes theoretical chemical principles in a Siamese neural network (14). By directly subtracting the latent spaces between the two component models, the authors model the relative free binding energy ΔΔ*G* between two ligands, enforcing the symmetric properties of relative binding affinities ΔΔ*G*_*AB*_ = Δ*G*_*A*_ − Δ*G*_*B*_ = −(Δ*G*_*B*_ − Δ*G*_*A*_). Active learning is also often used to incorporate experimental data into machine learning training procedures, guiding machine learning experiments with real-life data (15–17).

In the same spirit, AF2’s authors incorporate physiochemical constraints into their work by training with a “violation loss.” The violation loss is a component of AF2’s composite loss function that penalizes predictions for generating structures with overlapping (i.e., clashing) atoms or bond lengths and angles that deviate from the corresponding values in the literature. Despite the addition of the violation loss, AF2 predictions have significant stereochemical violations. The recommended prediction procedure involves a relaxation step with molecular dynamics force fields (MD FFs), which takes several seconds but significantly improves the final structure by removing many of these violations. Triangle point attention, also developed by AF2’s authors, is suggested to provide physical and geometric intuition to AF2. However, because it is a neural network component with learned parameters, it is unclear how it works in practice or if it genuinely imbues the model with any physical constraints.

## METHODS

This section describes our method to improve structure predictions from methods like AF2 by incorporating physical principals from molecular dynamics force fields. We will discuss the intuition for our approach, practical considerations for its development, and our procedure for training and evaluating our models with custom datasets.

OpenFold is an implementation of AlphaFold2 that attempts to faithfully reproduce AF2 in PyTorch with added utilities for training. OpenFold was developed by Ahdritz et al. (18), and AlphaFold2 by Jumper et al. (1). We refer to OpenFold and AlphaFold2 interchangeably as AF2, though we only work with the OpenFold code base.

### Forces from molecular dynamics can be interpreted as gradients in backpropagation

To improve overall protein structure prediction accuracy and to help protein structure prediction models better understand physical principles, we propose capitalizing on the relationship between the physical forces computed by molecular dynamics (MD) software (**Eq 1**) and the gradients utilized in machine learning backpropagation (**Eq 2**).

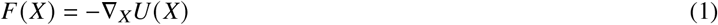

MD software evaluates the energy *U*(*X*) of a molecular system at each time step based on the positions of the atoms *X* and the force field parameters. Next, the software computes the forces acting on the atoms via **Eq 1**. Since MD software can readily compute *U*(*X*) and −∇_*X*_*U*(*X*) for a protein system, we propose treating *U*(*X*) as a loss function whose gradients are −*F*(*X*).

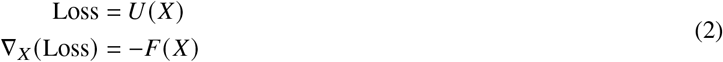

For backpropagation machinery to utilize these gradients during training, all that is needed is a custom layer with “forward” and “backward” components that return *U*(*X*) and −*F*(*X*), respectively. We can then apply this framework to any machine learning model capable of predicting all-atom protein structure representations.

### OpenMM-based loss function

Our custom loss function implements **Eq 2** via the steps depicted in **Fig 1. (1)** A deep learning model predicts an all-atom protein structure representation. Our code supports Cartesian and internal coordinate (bond and torsional angle) representations of protein structure. **(2)** The user determines a mapping between the atoms or angles predicted by their model and the atom or angle representation expected by the SidechainNet protein representation. SidechainNet (19) is a Python library and dataset with tools for PyTorch-based (20) deep learning protein structure prediction tasks. **(3)** In preparation for energy evaluation with OpenMM (21), a molecular dynamics system is initialized using the all-atom protein representation, and hydrogen atoms are added (if not already present) to the prediction using differentiable vector operations. **(4)** The custom loss function layer and SidechainNet protein representation compute a loss equal to the predicted structure’s potential energy. The forces acting on the atoms, also calculated by OpenMM, are designated as gradients to train the model. The model parameters are updated, and training continues.

**Figure 1:**
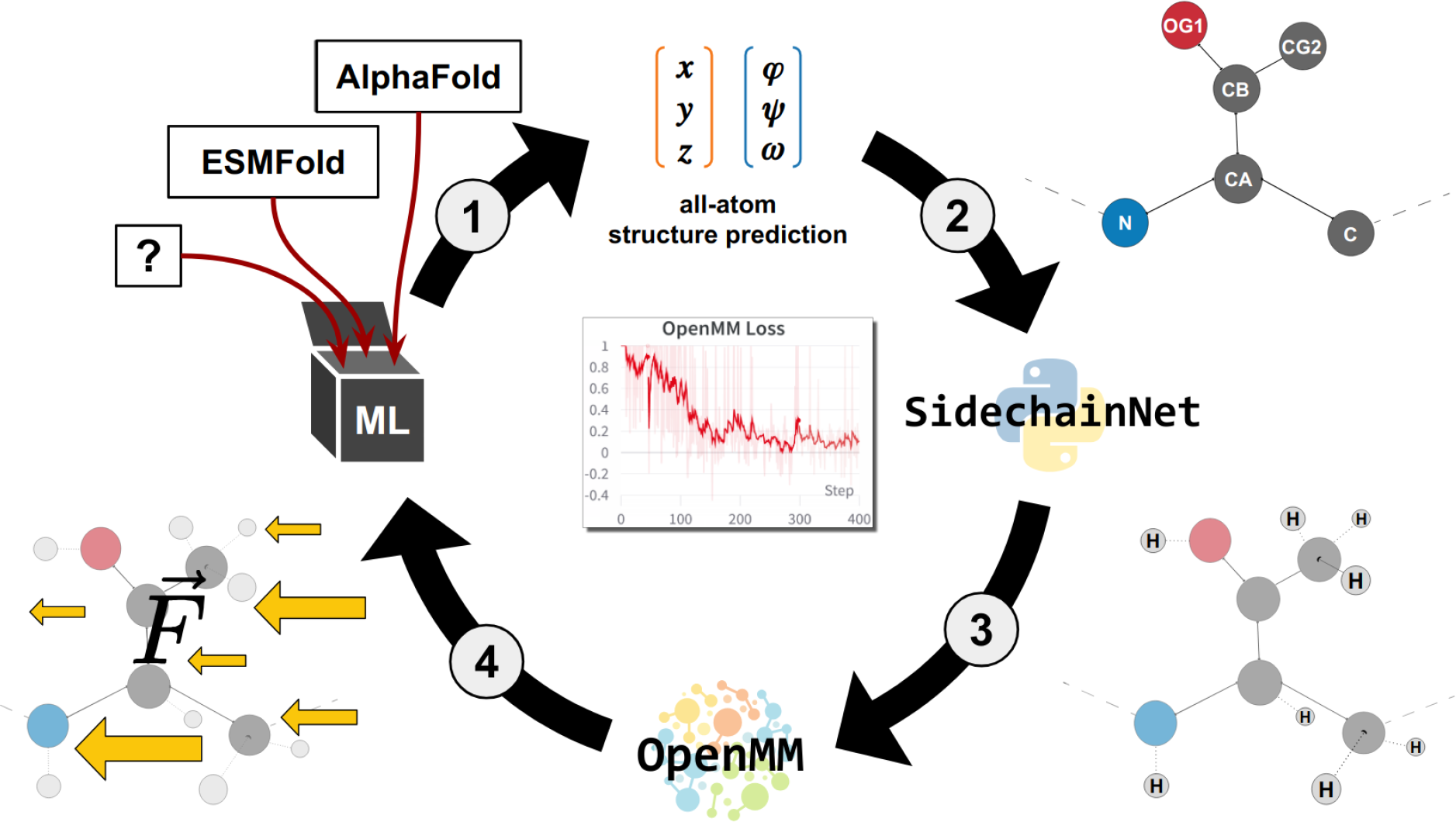
A graphical summary of our method for training deep learning protein structure prediction models by interpreting MD forces as gradients.

**Figure 2:**
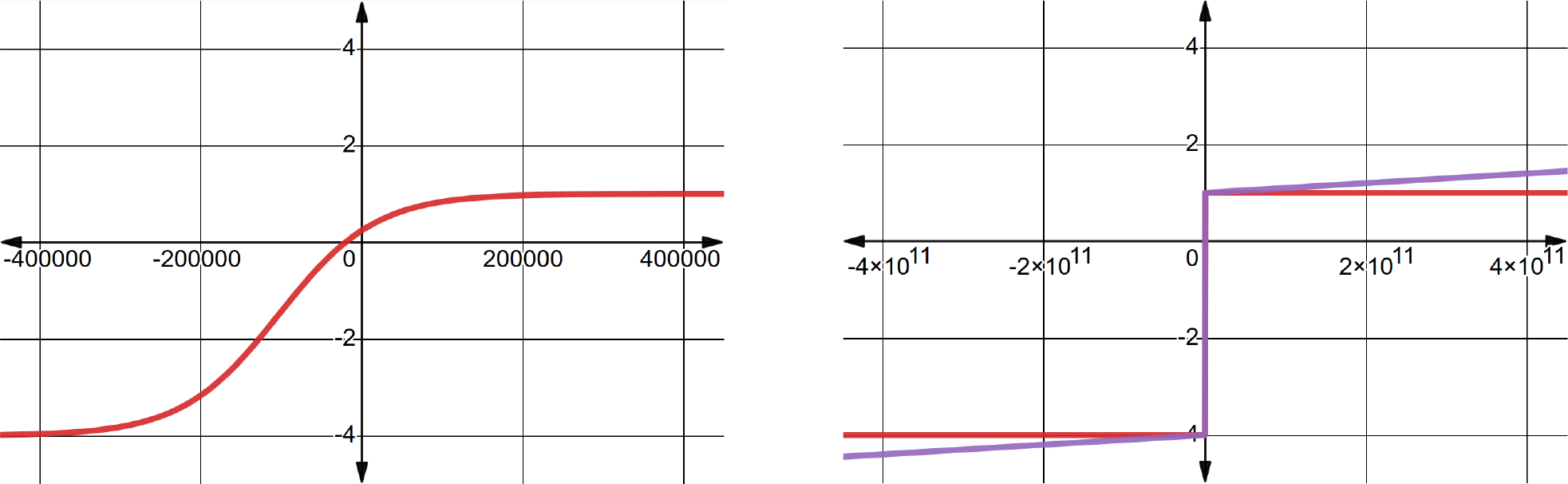
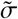 and σ^∗^ in red and purple, respectively. Both functions have nearly identical behavior, except σ^∗^ has a non-zero derivative for extreme values of *x* (energy).

Users may specify any force field and solvent model available in the OpenMM package for use with our method. For experiments in this paper, we selected the Generalized Born Implicit Solvent (`gbn`) model along with the `ff15ipq` force field (22).

#### Practical Consideration I: Hydrogens

Let *X*_*H*_ and 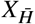 represent a predicted protein structure with and without hydrogen atoms. MD FFs require protein systems to contain hydrogen atoms in order to compute *U* (*X*_*H*_) and *F* (*X*_*H*_). However, AF2 and related models predict structures without hydrogens, 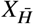. Thus, we must find a way to add hydrogens to the structures predicted by these models during training so that we may utilize the values computed by OpenMM. If hydrogens are added to the predicted structure in a non-differentiable manner (e.g., via OpenMM), we cannot train our model end-to-end. This is because we must be able to compute the gradients or construct a forward and backward component for every layer of our neural network. Adding hydrogens outside of PyTorch would not retain the computation graph necessary for automatic differentiation. In other words, we would not easily be able to compute the derivative of the function, *g*, that adds hydrogens to the predicted structure, 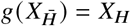.

We developed software in the SidechainNet package to address this issue. Our method adds hydrogens (if needed) to predicted protein structures, using only differentiable PyTorch vector operations to maintain differentiability. The software components found in SidechainNet’s `SCNProtein` class enable building hydrogen-inclusive Cartesian coordinates starting from angle-only representations via a sidechain-parallel method based on Natural extension of Reference Frame (NeRF) (23) or by adding hydrogens to the Cartesian coordinates of existing heavy atoms, also via NeRF. Next, our software sets up an OpenMM MD system. Our custom loss function is then used to compute *U*(*X*_*H*_) in the forward pass and −*F*(*X*_*H*_) in the backward pass by querying these values from OpenMM. Since converting 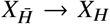 via vector operations is differentiable, PyTorch can effectively map the all-atom forces back to the heavy atom coordinate representation.

Our method adds hydrogens in consistent, predetermined conformations without further optimization. Although this placement may result in non-optimal hydrogen positions, this does not prevent our system from minimizing the structure’s potential energy.

#### Practical Consideration II: Rugged Potential Energy Landscapes

Training a machine learning model using potential energy as a loss function is challenging because protein energy landscapes can be incredibly rugged (24, 25). Small changes in sidechain rotamers may cause unrealistic clashes in otherwise perfect structures. Suppose MD FFs evaluate the energy of a protein system with clashes. In that case, the software may return astronomically high energy values (infinite or approaching the maximum value of computational numerical representation). These values reflect the physical impossibility of two atoms occupying the same physical space. This behavior is not frequently observed in MD trajectories because atoms often do not move enough between time steps to overlap or take on unrealistic conformations. However, when training DL models using mini-batches for gradient descent, each subsequent prediction is for a different protein and represents a new MD system. Thus, there are no guarantees that the predicted system will be in a realistic conformation. Furthermore, in a model like AF2, there are several other loss terms in the composite loss function to consider (e.g., Frame Aligned Point Error (FAPE) loss, distogram loss, and masked multiple sequence alignment loss). The raw energy of the predicted system would easily dominate the magnitude of the other loss terms, which we observed having maximal values of about 1.5 loss units in a trained AF2 model.

To smooth out the energy landscape’s ruggedness and transform the loss’s range into a range that is less likely to dominate other loss terms, we construct a sigmoid function, 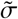, by modifying the standard sigmoid function, σ, and applying it directly to the raw energy, *x*.

| Parameter | Value |
| --- | --- |
| $A$ | 5 |
| $B$ | $10^6$ |
| $C$ | $3 \cdot 10^5$ |
| $D$ | 5 |

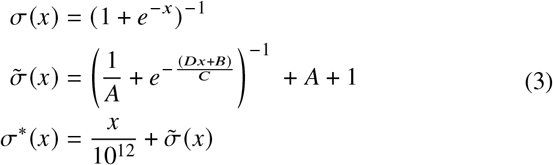

We heuristically chose several constant scalar parameters to construct a sigmoid function with desirable properties. 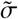 has a maximum value of 1, a minimum value of -4, and changes from positive to negative when the energy of a protein structure is about -20,000 kJ/mol. We also selected a loss component weight of 0.01 for the final computed loss value. These values were chosen so that the range of our loss function during training would not differ significantly from the range of values of the other loss components. For example, we reasoned that since AF2 loss components did not become negative during our finetuning experiments, we would like our energy-based loss to become negative only after surpassing a low-energy threshold of -20,000 kJ/mol that we associated with energetically favorable protein structures. Allowing our loss function to become negative would enable the model to optimize the predicted structure’s energy further, even when other loss components reached their minimal values.

We developed a second modified sigmoid, σ^∗^, which is a “leaky” version of 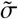. σ^∗^ adds an extremely small fraction of *x* to 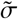. This enables constant, non-zero gradients for even extreme energy loss values. Theoretically, this allows the model to learn from physically implausible structures while still dramatically smoothing the energy-loss landscape. This modification also removes the asymptotic lower bound in 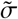 so that models may reach any energetic minimum.

In practice, we found these modifications helpful for training and visualizing energy values since the range of raw energy values of predictions can be too large to plot even on a log-scaled axis.

### Training

#### Models

To measure the effects of our custom loss function, we developed a fork of the OpenFold model that enables training with our data and loss function. We finetuned models rather than training from scratch, loading weights provided by the OpenFold authors.

We attempted to match the finetuning procedures described in the OpenFold and AF2 papers. However, we did not increase the Multiple Sequence Alignment (MSA) depth or crop size to limit GPU memory consumption. We trained each model using four A100s for about 12 days and achieved an effective batch size of 128 by accumulating gradients 32 times. To avoid training instability, we used an OpenMM-Loss-specific learning rate schedule that linearly scaled the loss function component weight to its maximum value over 1000 steps.

#### Data

We propose two distinct and mutually compatible ways to train AF2 to predict structures with lower energy and better physical characteristics: training on energetically minimized protein structures and training with our loss function.

We propose training on energetically minimized protein structures for the following reason. Suppose we train AF2 on raw, unminimized PDB structures with our loss function. As mentioned before, structures in the PDB likely do not reflect realistic structures in an aqueous environment. In that case, the accuracy-based (e.g., FAPE) and energy-based (e.g., OpenMM-Loss) loss function components may push the model in opposing directions, one towards accurately reflecting the PDB structure and another towards predicting a conformation with low energy. Training on protein structures that already exist at a local minimum of our loss function should enable the model to satisfy both objectives simultaneously.

To create this minimized dataset, we used our loss function (without the sigmoid component) to minimize a set of proteins from the SidechainNet dataset. We chose to minimize the energy of our proteins with respect to their bond and torsional angles rather than their atomic coordinates. This theoretically allows models that predict angle representations of proteins to predict their target with perfect accuracy since bond lengths and angles in the minimized set are set to idealized values with respect to the `ff15ipq` force field. Starting with about 100,000 protein entries from the SidechainNet CASP 12-era (26) dataset, we ended up with about 32,000 entries that (1) did not have gaps in their structures (a requirement for our minimization procedure); and (2) did not deviate more than 5 RMSD from their starting structure after minimization. This dataset is called **scnmin**.

An unminimized version of this dataset containing identical protein entries, called **scnunmin**, was retained to evaluate our loss function without minimized protein structures.

We utilized MSAs and template hits generated by the OpenFold authors for training when available (18). We followed the `mmseqs2`-based (27) procedure suggested by the OpenFold authors to generate the remaining input MSAs and template hits.

We chose a validation set from CAMEO (28) to match the one in the OpenFold paper. We constructed a new test set containing 93 proteins from a one-year window of CAMEO proteins ending on January 3, 2023. Our validation and test sets were minimized similarly to the training set while retaining their unminimized counterparts. In cases where a protein was rejected from our minimization procedure, we excluded it from both the minimized and unminimized sets. To determine training convergence, we used the minimized version of the validation set when training with **scnmin** and the unminimized validation set when training with **scnunmin**.

### Evaluation

It is essential to briefly discuss the distinction between accuracy (e.g., lDDT (29)) and structural quality (e.g., MolProbity Score). Methods papers for protein structure prediction have understandably focused on accurately predicting the protein structures available in the PDB. However, as discussed earlier, these structures are not ideal prediction objectives because they are potentially dissimilar from their *in vivo* counterparts. Additionally, their structures may change significantly in different environments or when bound to other ligands.

Accuracy metrics necessitate a selection of ground truth structures. Because we trained models to predict either minimized or unminimized structures from the PDB, using accuracy alone to assess our method’s performance would be insufficient. Instead, we propose that a more appropriate assessment of our method would be to compare orthogonal measures of structural quality—such as the MolProbity Score or the Clash Score computed by the MolProbity software suite (30). The MolProbity Score for a given structure reflects the crystallographic resolution at which the observed amount of structural irregularities would be expected. For example, a MolProbity Score of 0.8 would be the expected score from a structure determined from an X-ray crystallographic experiment with a resolution of 0.8 Å. The Clash Score measures per-contact clash energy in a protein structure when atoms are unrealistically close to each other. In both cases, lower values are better. We assess structural quality metrics before and after AF2-prescribed relaxation with OpenMM and AMBER force fields.

We repeated each finetuning experiment with three different random seeds to measure the robustness of our method. In the case of AF2, we utilized three different sets of weights reported to have similar performance instead of retraining AF2 from scratch (`finetuning_3.pt, finetuning_4.pt, and finetuning_5.pt`). In the subsequent plots and analysis, we combined the results from all three seeds for simplicity. However, the trends we observe are consistent across all seeds.

We checkpointed models by their performance on the validation set with respect to the OpenMM-Loss. We computed p-values using the one-sided T-Test implemented in `scipy` (31) with respect to the AF2 baseline.

## RESULTS

### OpenMM-Loss lowers prediction potential energy while maintaining accuracy

When trained with our loss function, predictions exhibit significantly lower energy loss values compared to predictions made from AF2 (**Fig 3**). Although we do not plot raw energy values (kJ/mol) here, we argue OpenMM-Loss is an appropriate substitution because it monotonically increases with energy and simplifies the interpretation of extreme values. lDDT_AA_, an all-atom accuracy metric, is not significantly different (p=0.8) for both methods when evaluated on the energy-minimized CAMEO test set (**Fig 4**). We use a simplified implementation of lDDT_AA_ that avoids renaming ambiguous atom names and thus slightly under-reports accuracy.

**Figure 3:**
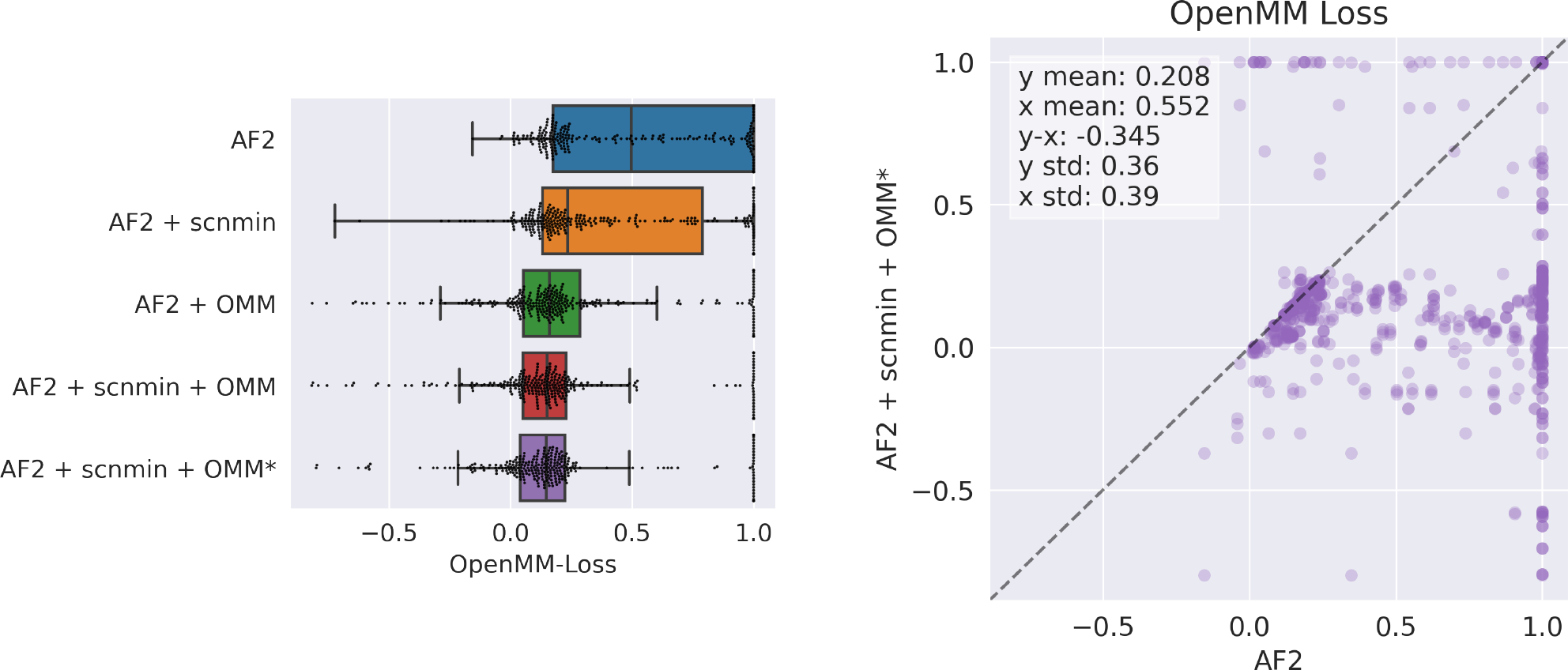
Model prediction potential energy represented by OpenMM-Loss. Each dot represents a predicted protein structure from the CAMEO test set before relaxation. **AF2**: baseline AlphaFold2 initialized using `finetuning_5.pt`. **+ scnmin**: finetuned on minimized SidechainNet proteins instead of unminimized structures. **+ OMM**: finetuned using OpenMM-Loss with 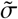 activation. **+ OMM**^∗^: finetuned using OpenMM-Loss with σ^∗^ activation.

**Figure 4:**
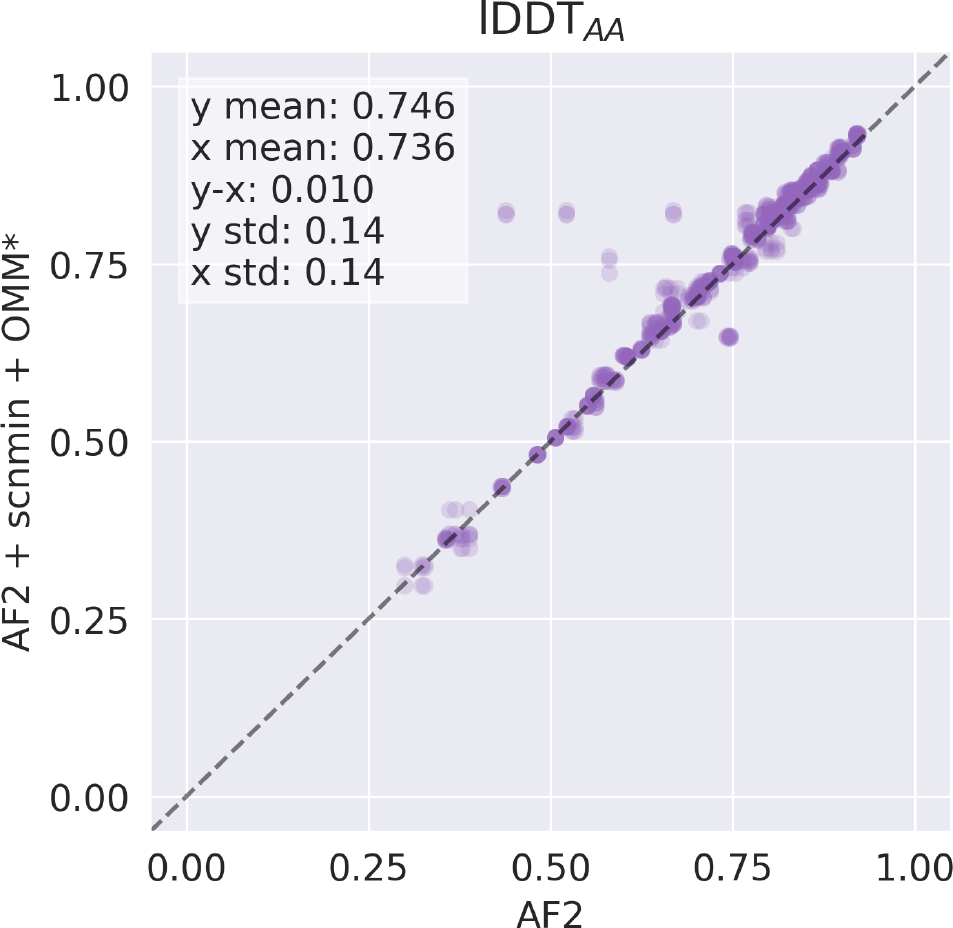
Accuracy between unrelaxed model predictions and the minimized CAMEO test when comparing our method and the **AF2** baseline. **AF2**: baseline AlphaFold2 initialized using `finetuning_5.pt`. **+ scnmin**: finetuned on minimized SidechainNet proteins instead of unminimized structures. **+ OMM**^∗^: finetuned using OpenMM-Loss with σ^∗^ activation.

### OpenMM-Loss improves physical properties of predictions before and after relaxation

Our method’s predictions have more favorable MolProbity Scores and Clash Scores on average than predictions from AF2 (**Fig 5, Fig 6**, and **Table 1**). Our method also has significantly more predictions with the lowest possible values of either metric (0.5 and 0, respectively). We observe a trend from top to bottom in **Fig 5** of improving structure quality as we add the components of our method (training on minimized structures, **scnmin**, and training with our custom loss, **OMM/OMM**^∗^).

**Table 1:** Mean structure quality metrics of structure predictions before and after relaxation. Computed p-values reflect the significance of being lower than the AF2 baseline.

| Model | MolProbity Score (p-value) |  | Clash Score (p-value) |  |
| --- | --- | --- | --- | --- |
|  | Unrelaxed | Relaxed | Unrelaxed | Relaxed |
| AF2 | 2.389 | 1.077 | 33.787 | 1.221 |
| AF2 + scnmin | 2.375 (4e-01) | 1.087 (6e-01) | 32.910 (3e-01) | 1.186 (4e-01) |
| AF2 + OMM | 2.285 (2e-02) | 0.986 (1e-02) | 28.156 (3e-06) | 0.744 (3e-06) |
| AF2 + scnmin + OMM | 2.271 (8e-03) | 0.945 (8e-04) | 27.143 (6e-08) | 0.598 (2e-09) |
| AF2 + scnmin + OMM* | **2.244** (1e-03) | **0.915** (3e-05) | **26.514** (2e-09) | **0.533** (2e-12) |

**Figure 5:**
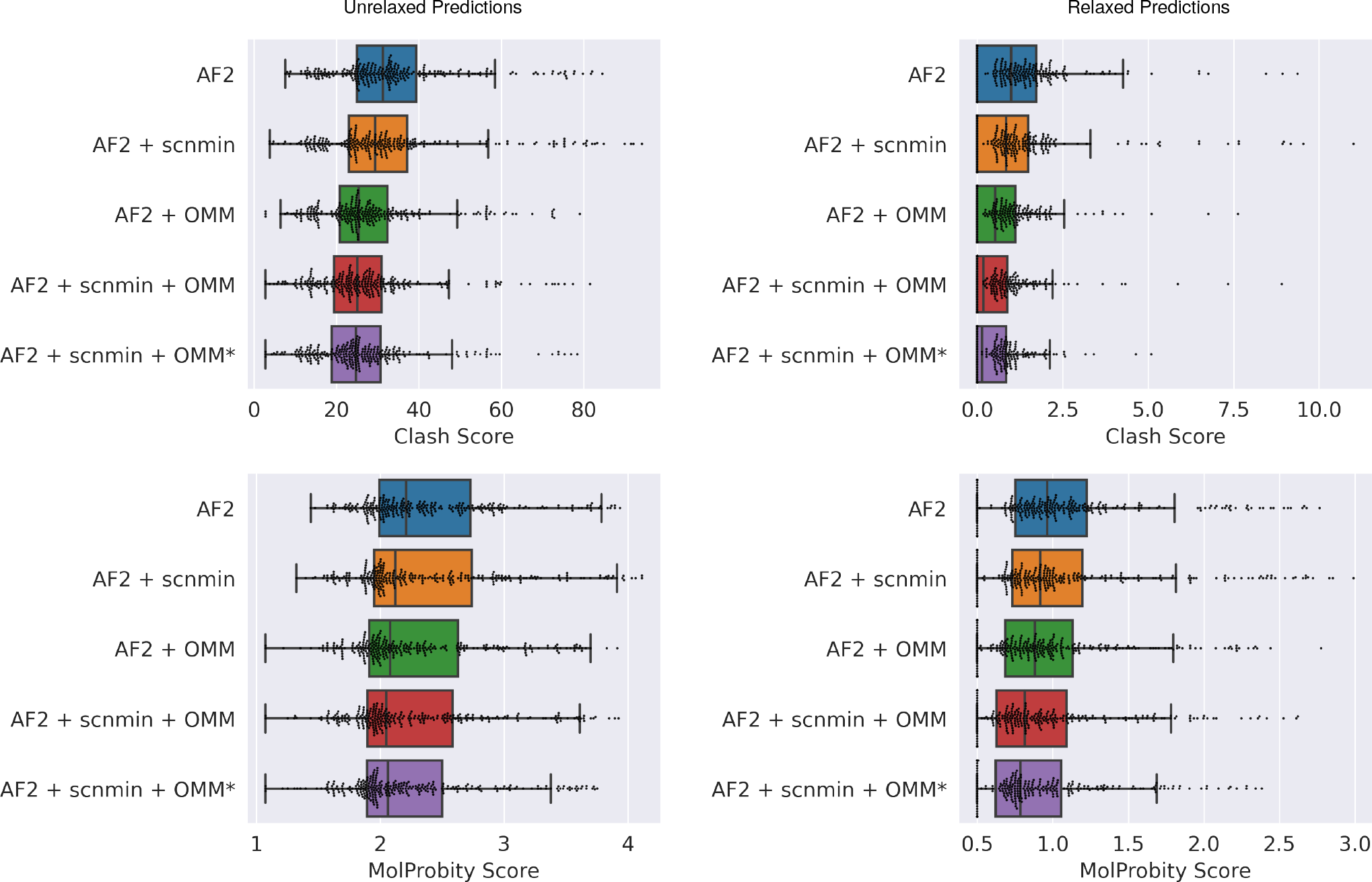
Evaluating the impact of the components of our method on structural quality metrics (Clash Score and MolProbity Score) relative to the **AF2** baseline. Each dot represents a single predicted protein from the CAMEO test set. Predictions are evaluated before and after relaxation (left and right columns, respectively). **AF2**: baseline AlphaFold2 initialized using `finetuning_5.pt`. **+ scnmin**: finetuned on minimized SidechainNet proteins instead of unminimized structures. **+ OMM**: finetuned using OpenMM-Loss with 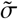 activation. **+ OMM**^∗^: finetuned using OpenMM-Loss with σ^∗^ activation.

**Figure 6:**
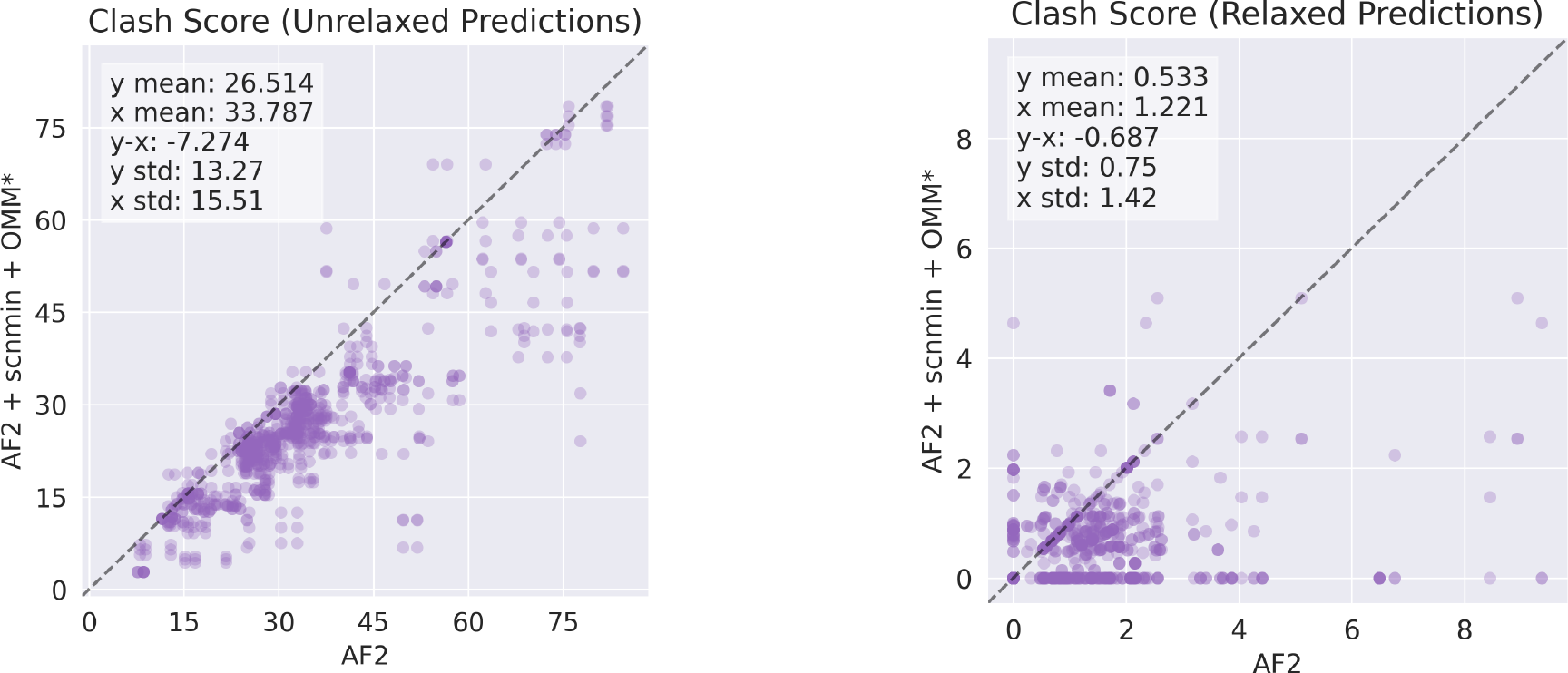
Comparing Clash Scores for predicted CAMEO test set proteins from one of our trained models (**AF2 + scnmin + OMM**^∗^) versus the **AF2** baseline. Model predictions are evaluated before and after relaxation (left and right columns, respectively). **AF2**: baseline AlphaFold2 initialized using `finetuning_5.pt`. **+ scnmin**: finetuned on minimized SidechainNet proteins instead of unminimized structures. **+ OMM**^∗^: finetuned using OpenMM-Loss with σ^∗^ activation.

It is worth noting that training on minimized structures (**scnmin**) alone does not significantly improve accuracy or structural quality metrics despite lowering the potential energy of predicted structures on average. In contrast, our loss function significantly improves structural quality relative to the **AF2** baseline. The best improvement is seen by combining these two training strategies.

The accuracy of our models between unrelaxed structures and the minimized CAMEO test set labels is shown in **Table 2**. Our objective is to produce more realistic predictions while not necessarily reproducing structures from the PDB, so we do not place a significant emphasis on accuracy values. It should be noted that direct comparison of accuracy metrics from the minimized CAMEO test set against the baseline AF2 has a caveat: AF2 is not trained to predict minimized structures, whereas all our methods are. Similar to our results regarding structural quality, we find that finetuning on minimized data and with OpenMM-Loss has a larger increase in accuracy than utilizing either one of these approaches individually.

**Table 2:**
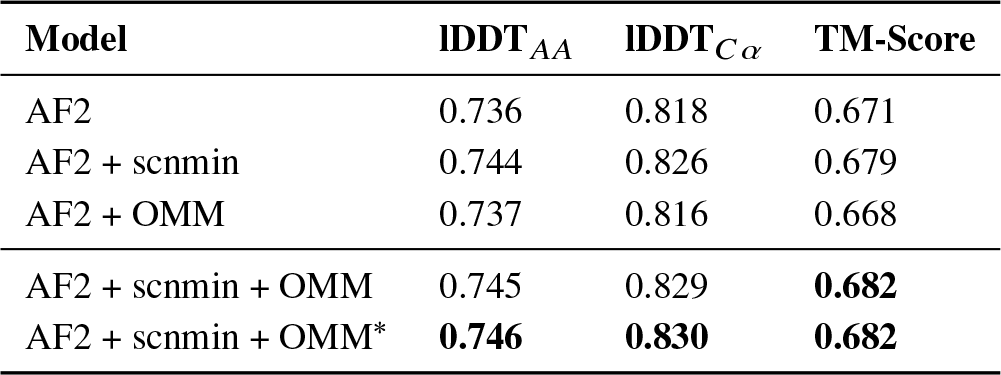
Structure prediction accuracy between unrelaxed predictions and the minimized CAMEO test set structures.

## DISCUSSION

If protein structure predictions are to become a valuable and practical tool for downstream tasks, the scientific community requires two things. First, we must improve accuracy at the sidechain level to avoid subtle changes in protein structure that have outsized impacts on tasks like molecular docking. Second, we require new paradigms to incorporate theoretical physical principles. By utilizing physical principles during training, it may be possible to continue the steady march towards improved accuracy while enforcing that predicted structures, at a minimum, pass the basic test of physical plausibility.

Accuracy in this experiment is difficult to define because we have compared models trained with minimized and unminimized labels. Still, we provide a framework for improving other important metrics related to the structural quality of protein structure predictions. We also hope to instigate a more thorough discussion on the relationship between structural accuracy (e.g., lDDT) and quality (e.g., MolProbity) metrics which are not always congruent.

A major limiting factor of current protein structure prediction methods is the lack of ligand information at training or evaluation time. A loss function like ours could help models better discriminate between favorable and unfavorable conformations of proteins and protein-ligand complexes by evaluating the potential energy of predicted systems during training. A model capable of understanding even a modest level of chemical interaction principles may have a significant advantage over models agnostic to such principles. Though our framework only supports protein representations at present, future work may study the impact of using an energy-based loss function for models that include protein-ligand or ligand-only systems.

## AUTHOR CONTRIBUTIONS

Jonathan King is a Ph.D. candidate in the research group led by David Koes. King and Koes designed the project’s goals and scope. King implemented the methods, carried out the experiments, performed the analysis, and wrote the manuscript.

## ACKNOWLEDGMENTS

This work is supported by R35GM140753 from the National Institute of General Medical Sciences, by NIH T32 training grant T32 EB009403 as part of the HHMI-NIBIB Interfaces Initiative, and in part by the University of Pittsburgh Center for Research Computing, RRID:SCR_022735, through the resources provided. Specifically, this work used the H2P cluster, which is supported by NSF award number OAC-2117681. The authors thank Andrew McNutt, Paul Francoeur, Rishal Aggarwal, and Katya Abelsky for their helpful comments during the preparation of this manuscript.

## SUPPLEMENTARY MATERIAL

Our fork of OpenFold that enables training with OpenMM-Loss can be found at https://github.com/jonathanking/openfold under a permissive Apache 2.0 License. Our custom loss function, along with supporting software and datasets, can be found at https://github.com/jonathanking/sidechainnet under a permissive BSD-3-Clause license.

